# Adiponectin plays a key role in macrophage-driven inflammation of adipose tissue caused by Chronic Intermittent Hypoxia

**DOI:** 10.1101/2025.09.10.675291

**Authors:** Xiaoqin Weng, Wenjing Wang, Mao Huang

## Abstract

**Background:** The inflammatory response driven by macrophages in adipose tissue is a key contributor to CIH-induced metabolic disorders. The precise mechanism underlying CIH-induced inflammation in adipose tissue macrophages remains elusive. Adiponectin (Ad) is a highly abundant adipokine secreted by adipocytes that plays a crucial role in insulin sensitivity.

**Methods:** This study aims to elucidate the pivotal roles of adipocytes and Ad in CIH-induced macrophage inflammation using a co-culture cell approach. The direct co-culture method involved the physical interaction between iBMDM cells and 3T3-L1 adipocytes, while the indirect co-culture method utilized adipocyte-conditioned media as the culture medium for macrophages. Following CIH treatment and Ad intervention, the expression levels of inflammatory factors in the conditioned media of each cell group were quantified. Statistical significance was defined as *P* < .05.

**Results:** Results showed CIH did not directly cause the release of inflammatory factors in individually cultured iBMDM cells and 3T3-L1 adipocytes; however, it significantly increased the release of inflammatory factors in co-cultured cells. Ad alleviates the pro-inflammatory effect of adipocytes on macrophages under CIH conditions.

**Conclusion:** CIH could disrupt adipocyte function and promote macrophage inflammation through paracrine mechanisms.

## 1. Introduction

Chronic Intermittent Hypoxia (CIH) is a key pathological feature of Obstructive Sleep Apnea (OSA) that has been identified as a major trigger for metabolic disorders. Clinical studies have shown a significant increase in CD68^+^ macrophage infiltration in the visceral adipose tissue of OSA patients, positively correlated with the severity of hypoxia.^1,2^ Animal models have further shown that 8 weeks of CIH exposure can increase the proportion of pro-inflammatory M1 macrophages in adipose tissue.^3^ Studies have shown that CIH activates TLR-4, leading to an inflammatory macrophage subset with high levels of IL-6 and IL-1β, which creates a chronic inflammatory environment in adipose tissue.^4^

This macrophage-driven inflammatory response disrupts metabolic homeostasis through multiple pathways: TNF-α secreted by adipose tissue activates the JNK pathway,^5^ leading to activation of Insulin Receptor Substrate 1 (IRS-1) and direct inhibition of insulin signaling.^6^ Meanwhile, IL-6 from macrophages activates the cAMP-PKA pathway in adipocytes, promoting triglyceride breakdown and the release of free fatty acids, which triggers hepatic lipid deposition.^7^ Notably, adipose-derived exosomes carrying inflammatory factors, such as miR-155, can remotely activate hepatic and endothelial inflammation, leading to a systemic cascade of inflammatory responses.^8^

The mechanism behind CIH-induced inflammation of adipose tissue macrophages remains unclear. An imbalance of adipokines is a key pathological factor in adipose tissue inflammation, with Adiponectin (Ad) being the most abundantly secreted insulin-sensitizing factor by adipocytes. Plasma levels of Adiponectin are significantly reduced in patients with OSA and negatively correlate with disease severity and hypoxia.^9,10^ Preclinical studies have shown that Ad binds to the AdipoR1 receptor on macrophage surfaces, activating the AMPKα signaling pathway to inhibit NF-κB nuclear translocation and STAT3 phosphorylation, ultimately decreasing the release of inflammatory factors such as TNF-α and IL-6, thereby impacting the stability of atherosclerotic plaques.^11^ In the respiratory field, evidence indicates a specific link between Ad levels and macrophage inflammation in asthma.^12,13^ Animal experiments also show that, in obese mice induced by a high-fat diet, exogenous administration of full-length Ad reduces macrophage infiltration into adipose tissue and restores the balance in M1/M2 macrophage proportions.^14^

Based on the evidence mentioned above, the main scientific debate centers on whether adipocyte involvement is essential and the role of Ad in CIH-induced inflammation of adipose tissue macrophages. Some in vitro studies support the direct-action hypothesis: isolated bone marrow-derived macrophages (BMDMs) can independently produce inflammatory responses under simulated CIH conditions (1% O_2_, 30 cycles/h).^4^ However, more in vivo studies reveal the existence of indirect regulatory mechanisms between adipocytes and macrophages. Research has shown that lactate produced by adipocytes can promote macrophage inflammation,^15^ exosomes derived from obese mice adipocytes induce M1 macrophage phenotypes by secreting miR-155,^8^ and Yepuri et al.^16^ have confirmed that in obese individuals, macrophages and adipocytes promote cardiac fibrosis through intercellular communication. This study aims to elucidate the vital roles of adipocytes and Ad in CIH-induced macrophage inflammation using a co-culture cell strategy.

## 2. Methods

### 2.1. Materials

High-glucose Dulbecco’s Modified Eagle Medium (DMEM) was obtained from Corning (New York, NY, USA). Both bovine serum and fetal bovine serum (FBS) were supplied by Gibco (California, USA). Enzyme-linked immunosorbent assay (ELISA) kits for detecting mouse TNF-α, IL-6, MCP-1, and IL-10 were purchased from Thermo Scientific (Rockford, IL, USA). The adipogenic differentiation medium for mouse bone marrow mesenchymal stem cells and Oil Red O stain were acquired from Procell Life Science (Wuhan, China). Ad (mouse) powder was sourced from Adipogen Life Sciences (Switzerland).

### 2.2. Cell culture and treatment

We used the immortalized bone marrow-derived macrophage (iBMDM) line (ATCC). Cells were cultured in DMEM medium with 10% heat-inactivated FBS at 37°C in a humidified incubator with 5% CO_2_.

Mouse 3T3-3L1 preadipocytes, also from ATCC, were differentiated using a commercial adipogenic induction medium mentioned earlier. In brief, cells were plated in 6-well plates and grown to full confluence. The medium was then replaced with an adipogenic differentiation fluid A for 48 hours, and then changed to a maintenance medium fluid B for 24 hours. After three rounds of induction by this pattern alternation, lipid accumulation was observed microscopically, and adipogenesis was confirmed via Oil Red O staining on day 9. Differentiated 3T3-L1 adipocytes were subsequently used for co-culture experiments.

### 2.3. Preparation of Ad culture medium

Ad was purchased from Adipogen in Switzerland, with a specification of 50 μg, and stored as a powder at −20℃. It was thawed and dissolved in serum-free DMEM to create a conditioned medium with a concentration of 10 μg/mL.

### 2.4. Coculture of adipocytes and iBMDM cells^17–19^

Adipocytes and iBMDM macrophages were co-cultured under both direct (physical contact) and indirect conditions. For direct co-culture, serum-starved adipocytes were seeded in 6-well plates, and iBMDM cells (5 × 10^5^) were added directly. In the indirect setup, conditioned medium collected from adipocyte cultures was applied to iBMDM cells. Cultures were maintained for 48 hours in serum-free DMEM with or without Ad supplementation. Monocultures of adipocytes and macrophages served as controls.

### 2.5. Cell CIH treatment^20^

The treated cells were cultured in serum-free DMEM medium within a three-gas cell culture incubator equipped with an oxygen (O_2_)/carbon dioxide (CO_2_)/nitrogen (N_2_) gas regulation system. The CO_2_ level was maintained at 5%, and the O_2_ level cyclically fluctuated between 1% and 21% every 15 minutes for a total of 30 minutes per cycle (15 minutes of low oxygen followed by 15 minutes of normal oxygen) over a continuous 48-hour period. Cell samples were then collected for subsequent analysis.

### 2.6. Cytokine measurement

Supernatants from both direct and indirect co-culture systems were collected after 48 hours.

Quantification of IL-1β, IL-6, TNF-α, MCP-1, and IL-10 levels was performed using commercially available ELISA kits (Thermo Scientific, Rockford, IL, USA) in strict accordance with the manufacturer’s instructions.

### 2.7. Statistical analysis

Results are shown as mean ± SEM from a minimum of three independent replicates. The choice of statistical test (Student’s t-test, one-way ANOVA with Tukey’s honest significant difference test, or Wilcoxon rank-sum test) was based on data distribution and the number of groups compared, after confirming assumptions of normality and equal variances. All analyses were performed using SPSS version 20, and a *P*-value below 0.05 was deemed statistically significant.

## 3. Results

1. CIH did not trigger the release of inflammatory factors in individually cultured iBMDM cells and 3T3-L1 adipocytes, but could promote the release of a large amount of inflammatory factors in co-cultured cells. To understand how macrophages contribute to CIH-related inflammatory responses in adipose tissue, we examined intercellular communication within a co-culture system of adipocytes and macrophages. Since the iBMDM cell line can mimic the phenotype of bone marrow-derived macrophages infiltrating adipose tissue under CIH conditions, we first investigated the effect of individual CIH treatments on the inflammatory response of macrophages and adipocytes. CIH interventions were applied separately to 3T3-L1 adipocytes and iBMDM macrophages. No significant increase in the secretion of inflammatory cytokines was observed in either cell type following individual CIH exposure; the concentrations of TNF-α, IL-6, and IL-1β were similar to those in the normoxic group (Figure 1a-1c), indicating that a response from a single cell type alone was not enough to trigger the inflammation cascade. Subsequently, we directly co-cultured adipocytes with iBMDM cells and exposed them to CIH treatment. Notably, co-culture alone significantly increased the expression of TNF-α in the cell culture supernatant, while CIH treatment further amplified the inflammatory response, resulting in a notable rise in the expression of TNF-α, IL-6, and IL-1β in the culture supernatant (Figure 1a-1c). These findings imply that physical contact between adipocytes and macrophages or paracrine signaling may have activated inflammatory pathways.
2. Macrophage inflammation in adipose tissue under CIH conditions is primarily elicited by paracrine signals originating from adipocytes. To understand the mechanisms of intercellular communication, iBMDMs were exposed to adipocyte-conditioned media after adipocytes were treated under normoxia or CIH. The results showed that conditioned media from normoxic adipocytes (iBMDM + AD-CM group) stimulated increased secretion of IL-6 and TNF-α by iBMDM cells. When treated with conditioned media from CIH-preconditioned adipocytes, a marked increase was observed in the secretion of TNF-α and IL-6 into the cell culture supernatant. (Figure 2a-2b). Although MCP-1 expression remained unchanged under normoxic conditions, it was significantly upregulated after CIH treatment (Figure 2c). IL-1β and IL-10 expression levels remained stable and showed no significant alteration (Figure 2d-2e). It can be inferred from these results that macrophage inflammation in adipose tissue induced by CIH is mainly driven by soluble factors secreted by adipocytes, which trigger macrophages to release inflammatory mediators, especially M1 pro-inflammatory cytokines.
3. Ad enhances the pro-inflammatory effect of adipocytes on macrophages under CIH exposure In previous studies, we found that adipocytes can promote macrophage inflammation through paracrine signaling. The conditioned media from adipocyte cultures after CIH treatment could induce macrophages to release more inflammatory factors such as TNF-α, IL-6, and the inflammatory chemokine MCP-1. We further analyzed the conditioned media from adipocyte cultures treated with and without CIH exposure. We observed a significant decrease in Ad production in adipocytes after CIH treatment, accompanied by an increased release of MCP-1 (Figure 3a-3b). Therefore, we hypothesize that the downregulation of Ad in adipocytes induced by CIH may lead to increased release of pro-inflammatory factors, which, through paracrine signaling, promote macrophage inflammation. Supplementation with Ad may improve the pro-inflammatory effect of adipocytes on macrophages under CIH exposure. By adding Ad to the culture medium of iBMDM, we observed that, compared to the iBMDM + CIH-AD-CM group, the levels of TNF-α (Figure 3c), IL-6 (Figure 3d), and MCP-1 (Figure 3e) in the conditioned media of macrophage cultures in the Ad intervention group (iBMDM + CIH-AD-CM + Ad) were significantly reduced. The expression levels of IL-10 (Figure 3f) and IL-1β (Figure 3g) showed no significant changes among the macrophage groups. These results indicate that Ad can effectively decrease the release of inflammatory factors and chemokines from macrophages induced by CIH, further confirming its role in improving adipocyte-mediated macrophage inflammation under CIH exposure.

**Figure 1.**
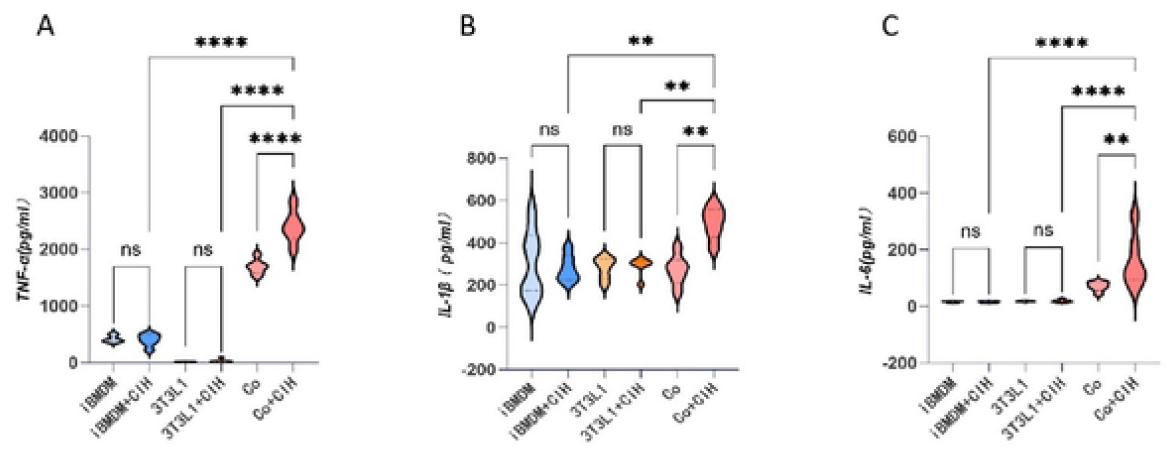
Secretion levels of the pro-inflammatory cytokines TNF-α (A), IL-1β (B), and IL-6 (C) in the cell culture supernatant after individual and combined co-cultures of iBMDM and 3T3-L1 adipocytes following CIH treatment. Values are expressed as mean ± SEM (n = 6). ^*^P < .05, ^**^P < .01, ^***^P < .001, ^****^P < .0001, with ns indicating not statistically significant.

**Figure 2.**
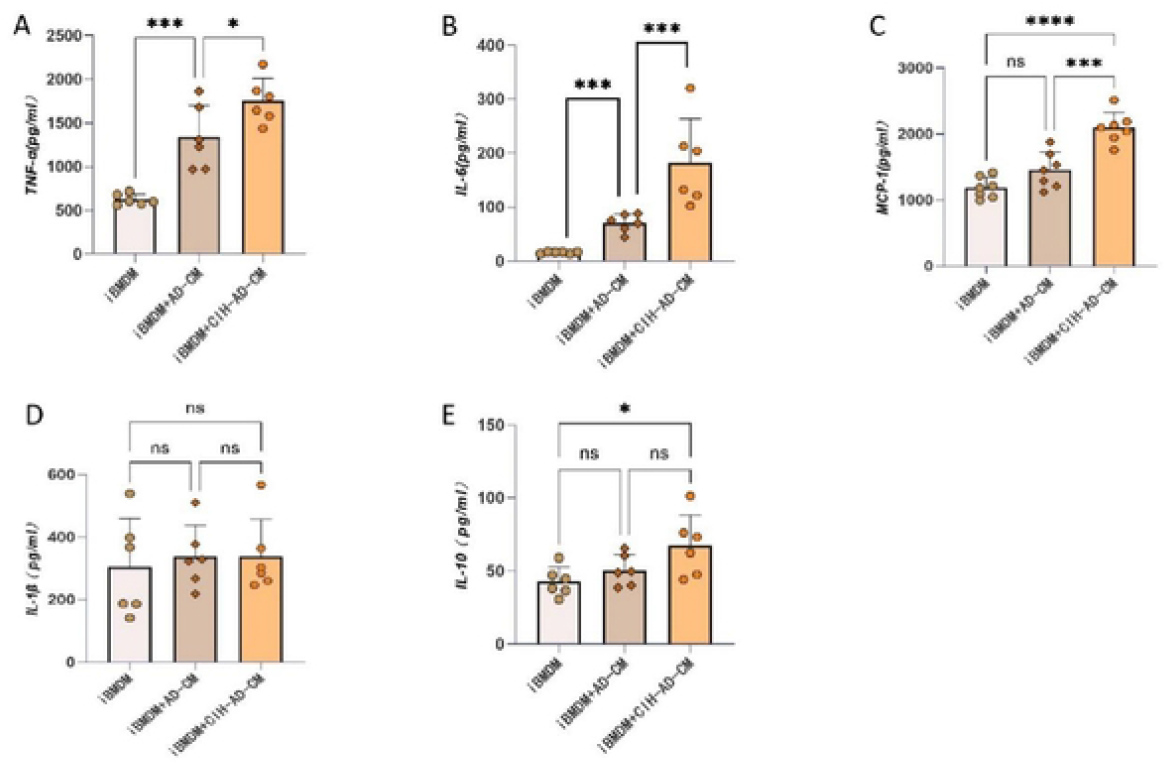
Cytokine secretion profile of iBMDM cells. The expression levels of TNF-α (A), IL-6 (B), MCP-1 (C), IL-1β (D), and IL-10 (E) were measured by ELISA in supernatants from iBMDM cells left untreated or stimulated with conditioned media from control (Ad-CM) or CIH-treated adipocytes (CIH-Ad-CM). Values are expressed as mean ± SEM (n = 6). ^*^P < .05, ^**^P < .01, ^***^P < .001, ^****^P < .0001, with ns indicating not statistically significant.

**Figure 3.**
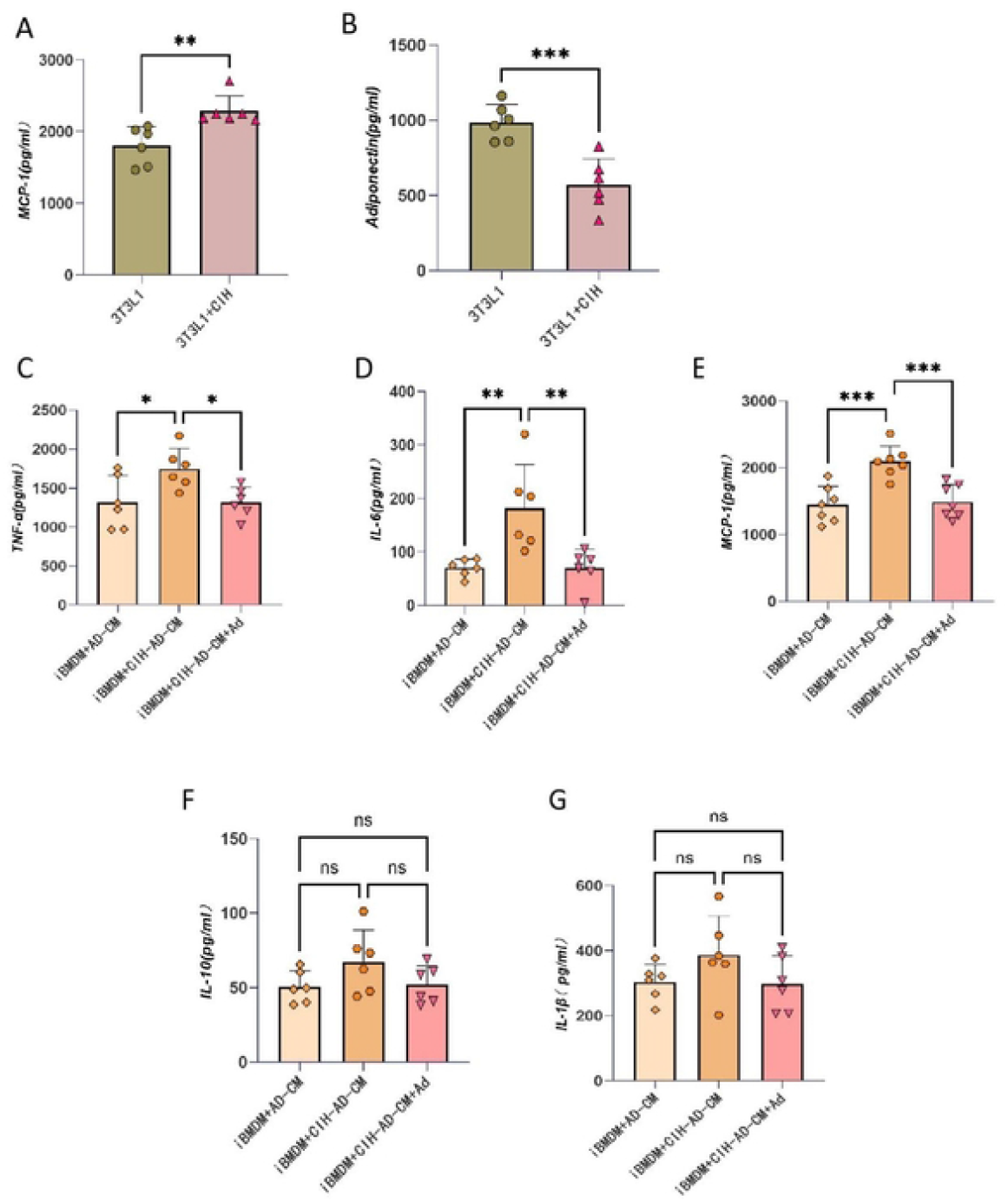
A-B. Expression levels of MCP-1 (A) and Ad (B) in the conditioned media of 3T3-L1 adipocyte cultures after CIH exposure; B. Levels of TNF-α (C), IL-6 (D), MCP-1 (E), IL-10 (F), and IL-1β (G) released in the conditioned media of macrophage cultures following Ad intervention. Values are expressed as mean ± SEM (n = 6). ^*^P < .05, ^**^P < .01, ^***^P < .001, ^****^P < .0001, with ns indicating not statistically significant.

## 4. Discussion

This study aimed to explore the sources and key regulatory mechanisms of macrophage inflammation in adipose tissue under CIH conditions. Our experimental results clearly show that CIH does not directly cause inflammation in macrophages but instead stimulates macrophage inflammation indirectly by affecting the paracrine function of adipocytes. In this process, the reduction in Ad secretion by adipocytes plays a vital role.

Firstly, our series of experiments successfully demonstrated that adipocytes are essential mediators of macrophage inflammation under CIH conditions. Results from the first part of the experiments showed that neither macrophages nor adipocytes exhibited significant inflammatory responses when individually exposed to CIH stimulation in vitro. This finding ruled out a direct pro-inflammatory effect of CIH on these two cell types. However, when the two cell types were co-cultured, CIH stimulation led to a significant upregulation of inflammatory factor expression. This strongly suggests that communication between the two cell types is a necessary condition for CIH-induced inflammation. Extending the observations by Wang et al.,^21^ who identified TNF-α upregulation in a 3T3-L1/RAW 264.7 co-culture system, our data further demonstrate that this pro-inflammatory response is mainly triggered by soluble factors secreted from adipocytes under CIH conditions. This confirms that adipocytes can promote macrophage inflammation by releasing soluble molecules and worsening adipose tissue inflammation.

To clarify the nature of this communication, we used an indirect co-culture system and found that adipocyte-conditioned media alone were enough to activate macrophages, with this effect being stronger under CIH stimulation. This clearly demonstrates that CIH influences the secretion of various soluble factors by adipocytes (i.e., paracrine effects), which then promote macrophage inflammation. It confirms that adipocyte-driven macrophage inflammation under CIH conditions is indeed mediated by adipocytes.

Furthermore, we explored the key factors involved in adipocyte paracrine effects, focusing on Ad. Under CIH stimulation, adipocytes not only secreted known inflammatory chemokines like MCP-1, but more importantly, the expression of the protective factor Ad significantly decreased. This offered a key mechanistic hypothesis: CIH may disrupt the balance of adipocyte secretion profiles, with an increase in pro-inflammatory signals such as MCP-1 and a decrease in anti-inflammatory signals like Ad happening simultaneously, collectively pushing macrophages toward an inflammatory phenotype. To confirm the functional role of reduced Ad, we performed rescue experiments, which showed that exogenous Ad effectively reversed macrophage inflammation induced by adipocyte-conditioned media. This outcome not only suggests that a decrease in Ad is necessary for inflammation but also directly demonstrates that the reduction of Ad is a primary cause of adipocyte-mediated macrophage inflammation under CIH conditions. Usually, Ad inhibits macrophage inflammation through its well-known anti-inflammatory pathways (such as activating AMPK). The decline in Ad caused by CIH removes this inhibition, making macrophages more susceptible to recruitment and activation by chemokines like MCP-1.

## 5. Conclusion

This study presents a new theory on how CIH causes adipose tissue inflammation: CIH disrupts adipocyte function and promotes macrophage inflammation through paracrine effects. The main limitation of this study is the incomplete understanding of other factors secreted by adipocytes and their combined effects with Ad. Additionally, more research at the molecular level is necessary to understand how specific signaling pathways downstream of Ad are controlled in the context of CIH. Future studies could focus on developing methods to increase Ad levels in fat tissue or replicate its effects, offering a promising target for preventing and treating OSA-related metabolic issues like insulin resistance.

## Abbreviations

Ad: Adiponectin
CIH: Chronic Intermittent Hypoxia
OSA: Obstructive Sleep Apnea
IRS-1: Insulin Receptor Substrate 1
BMDMs: bone marrow-derived macrophages
DMEM: Dulbecco’s Modified Eagle Medium
FBS: fetal bovine serum
ELISA: Enzyme-linked immunosorbent assay
iBMDM: immortalized bone marrow-derived macrophage
O_2_: oxygen
CO_2_: carbon dioxide
N_2_: nitrogen

